# Large scale and significant expression from pseudogenes in *Sodalis glossinidius* - a facultative bacterial endosymbiont

**DOI:** 10.1101/124388

**Authors:** Ian Goodhead, Frances Blow, Philip Brownridge, Margaret Hughes, John Kenny, Ritesh Krishna, Lynn McLean, Pisut Pongchaikul, Rob Beynon, Alistair C. Darby

## Abstract

The majority of bacterial genomes have high coding efficiencies, but there are some genomes of intracellular bacteria that have low gene density. The genome of the endosymbiont *Sodalis glossinidius* contains almost 50% pseudogenes containing mutations that putatively silence them at the genomic level. We have applied multiple ‘omic strategies, combining: Illumina and Pacific Biosciences Single-Molecule Real Time DNA-sequencing and annotation; stranded RNA-sequencing; and proteome analysis to better understand the transcriptional and translational landscape of *Sodalis* pseudogenes, and potential mechanisms for their control. Between 53% and 74% of the *Sodalis* transcriptome remains active in cell-free culture. Mean sense transcription from Coding Domain Sequences (CDS) is four-times greater than that from pseudogenes. Comparative genomic analysis of six Illumina-sequenced *Sodalis* isolates from different host Glossina species shows pseudogenes make up ~40% of the 2,729 genes in the core genome, suggesting that they are stable and/or *Sodalis* is a recent introduction across the *Glossina* genus as a facultative symbiont. These data further shed light on the importance of transcriptional and translational control in deciphering host-microbe interactions. The combination of genomics, transcriptomics and proteomics give a multidimensional perspective for studying prokaryotic genomes with a view to elucidating evolutionary adaptation to novel environmental niches.

**Importance:** Bacterial genes are generally 1Kb in length, organized efficiently (i.e. with few gaps between genes or operons), and few open reading frames (ORFs) lack any predicted function. Intracellular bacteria have been removed from extracellular selection pressures acting on pathways of declining importance to fitness and thus, these bacteria tend to delete redundant genes in favour of smaller functional repertoires. In the genomes of endosymbionts with a recent evolutionary relationship with their host, however, this process of genome reduction is not complete; Genes and pathways may be at an intermediate stage, undergoing mutation linked to reduced selection and small population numbers being vertically transmitted from mother to offspring in their hosts, resulting in an increase in abundance of pseudogenes and reduced coding capacities. A greater knowledge of the genomic architecture of persistent pseudogenes, with respect to their DNA structure, mRNA transcription and even putative translation to protein products, will lead to a better understanding of the evolutionary trajectory of endosymbiont genomes, many of which have important roles in arthropod ecology.

## Introduction

The genomes of intracellular parasites and endosymbiotic bacteria evolve under conditions that are fundamentally different to those of free-living organisms (1). In many arthropod systems, bacteria can provide nutrients that are otherwise scarce to their host (such as B-vitamins absent from blood meals, or essential amino acids absent from plant sap), in exchange for host provision of protection, nutrition and mechanisms for vertical or horizontal transmission (2, 3). Obligate intracellular symbionts are maintained by the host and have evolved strategies that ensure their vertical transmission to the next generation of hosts. Ultimately this intracellular lifestyle, small population size and strict vertical transmission can result in extremely reduced genomes (1, 3–6). The general theory and process of this extreme genome reduction has been well studied using genomic data for intracellular bacteria, including endosymbionts such as *Buchnera* in aphids (7, 8), and *Wigglesworthia* in tsetse flies (9). However, gene loss is not limited to obligate intracellular pathogen/symbionts with strict vertical transmission, it is also observed in free-living bacteria and facultative symbionts (10).

One of the most important mechanisms for gene loss is that of pseudogenisation, resulting from the accumulation of nonsense mutations in protein coding sequences (1). These mutations putatively silence the gene at the genomic level resulting in theoretically non-functional genes/proteins (11). Prokaryotic pseudogenes generally exist at levels approximately between 1% and 5% (12). Comparative genomic analysis between closely related strains suggests that pseudogenes are often associated with reduced selective pressure on redundant gene sets allowing mutation to accumulate and inactivate genes. This has been observed as *Salmonella* changes host range or utilizes a new environment (13). The low level of pseudogenes in most bacteria suggests that they are removed rapidly from the genomes due to strong selection for genome efficiency (11). There are however examples among the intracellular pathogens and endosymbionts of high levels of pseudogene presence, reducing coding capacity down towards 50% in *Sodalis glossinidius* (14) and *Mycobacterium leprae* (15). Likewise, pseudogenes can persist for long periods – the mean half-life of *Buchnera aphidocola* pseudogenes has been estimated to be 24 million years (16). Pseudogenes have been well studied in the context of comparative genomics to understand how gene loss has shaped bacterial genomes (17), but whether they continue to contribute to the genetic capabilities of the bacterium has seldom been assessed (18). It could, for instance, be suggested that if pseudogene-derived transcription retains some form of *cis/trans* regulatory function, then this could select for pseudogene retention in the genome (19). It is also clear that under some circumstances, specifically where polymerase infidelity corrects for a frameshift within homopolymeric tracts at the transcriptional level, pseudogenes can still produce functional proteins that contribute to the fitness of the bacterium (20).

In this study we aim to understand the importance of pseudogenes in bacterial genome evolution in a model of a degrading bacterial genome, that of *Sodalis glossinidius*. *Sodalis* is a facultative intracellular, secondary endosymbiont of the tsetse fly (Diptera: *Glossina*). The variable frequency of *Sodalis* in natural populations suggests that *Sodalis* is not an obligatory component of the tsetse microbiome (21), however, the occurrence of *Sodalis* in natural populations has been linked to an increased capacity of tsetse to vector African trypanosomes (22).

Interestingly, *Sodalis* has a relatively large genome for a facultatively intracellular endosymbiont (~4Mbp) and two genome annotations suggest that pseudogene levels are between 29% (14) and 38% (23) of the total gene content. The *S. glossinidius* genome has amongst its coding repertoire, systems for flagella, transmembrane transport (14), quorum sensing (24), and, of note, type-III secretion systems, encoded by three *Sodalis* symbiosis regions (SSR 1-3), which are analagous to pathogenicity islands (e.g. Salmonella Pathogenicity Islands (SPIs (25)) and have been implicated in establishing or maintaining symbiosis (26). By combining the latest high-throughput sequencing and proteomics methods, we hope to shed light on potential post-transcriptional regulatory mechanisms that may be mitigating any potential deleterious effects. At the RNA level, riboswitches (27), or small RNAs (sRNAs) – short, 50-300bp transcripts mediated by imperfect base pairing interactions, have been shown to regulate genes in this manner (28). DNA methylation could also serve as a mechanism by which to control transcription and/or translation (29, 30).

*Sodalis glossinidius* represents an ideal system in which to test hypotheses surrounding pseudogene functionality and their evolution, as the organism maintains an unusually reduced coding capacity, yet remains amenable to cell culture allowing for sufficient DNA, RNA and peptides to be extracted for poly-omic analyses. First: Assuming genes with nonsense mutations are non-functional and therefore costly to the cell, pseudogenes should be evolving rapidly and be removed from the genome. Secondly: If pseudogene transcription or translation is deleterious, pseudogenes should be transcriptionally and translationally silent. Thirdly: Given hypothesis two, we can expect there to be genetic mechanisms to silence pseudogenes, and we will be able to identify genetic and transcriptional features that determine pseudogene status using a combination of genomic, expression and proteomic analysis. To that end, we tested these hypotheses by: 1. Establishing pseudogene content and evolution using pan-genome data; 2. Evaluating genome-wide methylation data and negative strand expression to elucidate potential expression control mechanisms; and 3. Correlating mRNA and protein expression levels to understand functional control of pseudogenes.

## Results

### *Sodalis* Genomics

To provide an updated, accurate reference for transcriptome mapping of the *S. glossinidius* isolate used in this study, we sequenced *de novo* a *Sodalis glossinidius* (from the host *Glossina morsitans morsitans*) isolate (SgGMMB4) using Pacific Biosciences sequencing. A single SMRTcell produced a total of 48,519 reads, with a mean length of 10,290 bp and mean read score of 0.85. The chromosome was assembled in to a single 4.1 Mbp contig and one copy of the three plasmid sequences pSG1 (90.747 bp); pSG2 (38,394 bp) and pSG3 (10,640 bp). Previously published annotations from the GMM4 isolate (14, 23, 31) were used, alongside a manually curated PROKKA-generated annotation of the PacBio sequenced isolate, to generate a new annotation of our *Sodalis* sequence (32). The overall mean genome GC content is 54.4 %, and the pseudogenes and CDS have a similar GC of ~55.5%. The revised annotation presented contains 3,336 putative CDS and 2,286 putative pseudogenes. In addition, 43 putative riboswitch domains, belonging to seven families, were identified (Supplementary Table 2). A prodigal (v. 2.6.2) (33), gene prediction including a Ribosome Binding Site (RBS) identification scan suggested a prevalence of standard methionine-coding ATG start codons (78 %), which decreases in the case of pseudogenes (67 %). GTG start-codons are the next most common, found in 14 % of all genes, increasing to 21 % of the total number of pseudogenes); followed by TTG (8 % overall; 12 % of pseudogenes). Sixty-one percent of all genes (48 % of pseudogenes) have an RBS predicted within 5-10 bp of the start codon, and 26 % of all genes (38 % pseudogenes) have no discernible RBS. A mummer-plot (34) of plasmid pSG1 reveals a 6,371 bp tandem repeat encompassing the inactive Type IV secretion system operon that is not present in the original *Sodalis* sequencing and annotation experiments, perhaps due to a collapsed repeat missed by first-generation sequencing assembly. Pseudogenes tend to be enriched for activities such as: transmembrane transport, including metal ions (GO:0099132; 23 pseudogenes / 10 CDS; Fisher Exact Test p=0.001); transposition (GO:0006313; 52 pseudogenes / 14 CDS; p<0.0001); receptors (GO:0004872; 27 pseudogenes / 13 CDS; p<0.0001) and glycerol metabolic processes (GO:0006071; 6 pseudogenes, 0 CDS; p<0.01).

### *Sodalis* Transcriptomics

To ascertain whether pseudogenes are being transcribed, or if their transcription is being regulated throughout growth, stranded RNA-sequencing was performed on three-replicates in three conditions across a bacterial growth experiment in cell-free media (Early Log Phase, ELP; Late Log Phase, LLP; Late Stationary Phase, LSP). Figure 1A shows boxplots of mean sense and antisense transcription (log (transcripts per million + 1) for each condition. From the density plots of sense transcription (Figure 1B) and of overall transcription (Figure 1C), it can be seen that there is a clear signal of no transcription from putatively inactive genes (logTPM+1=0). Other studies have used transcripts per million (TPM) of ≥1 (35) or ≥10 (36) as an indicator of activity, which are displayed on each of these figures. For this study, we have defined putatively active genes as having an arbitrary TPM value ≥ 1 in all three biological replicates in at least one condition. Genes with TPM≥10 in all three biological replicates in at least one condition are additionally described as being active. 53% of all combined genes and pseudogenes, 3,087 (2,237 CDS; 850 pseudogenes) exhibit active sense transcription (TPM≥10) in any given condition, with an additional 1,191 (547 CDS; 644 pseudogenes) putatively active (TPM ≥ 1; A total of 73.7% of all genes and pseudogenes). Additionally, 1,088 genes (703 CDS; 385 pseudogenes) showed active antisense transcription, with an additional 993 genes (629 CDS; 364 pseudogenes) exhibiting putatively active antisense transcription according to the same rules.

**Figure 1A.**
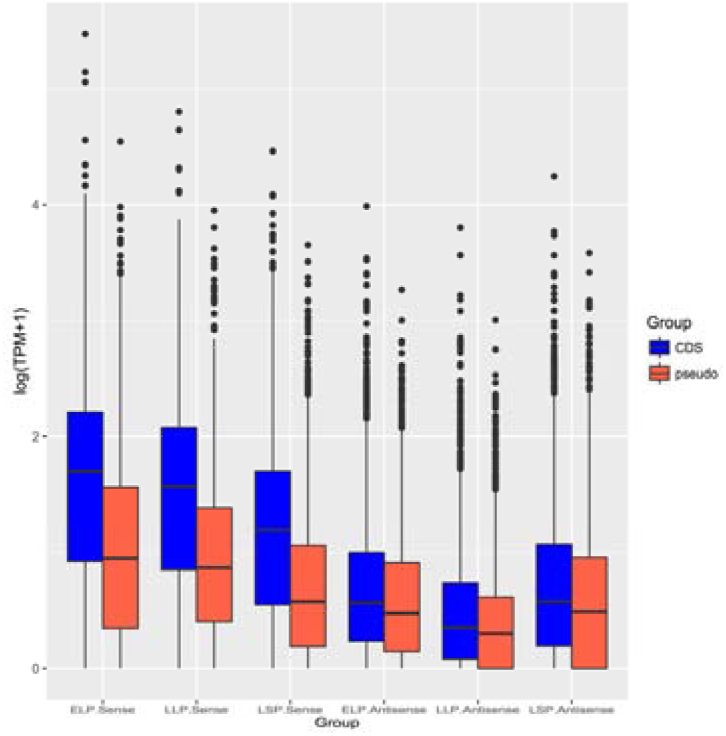
Sense and antisense mean transcription of CDS (blue) and pseudogenes (red) in three cell-free cultures of *Sodalis glossinidius*: Early Log Phase (ELP); Late Log Phase (LLP) and Late Stationary Phase (LSP). Transcripts per million (TPM) derived from EdgeR counts per million have been transformed to log(TPM+1) to enable presentation of zero transcription.

**Figure 1B (left) and 1C (right).**
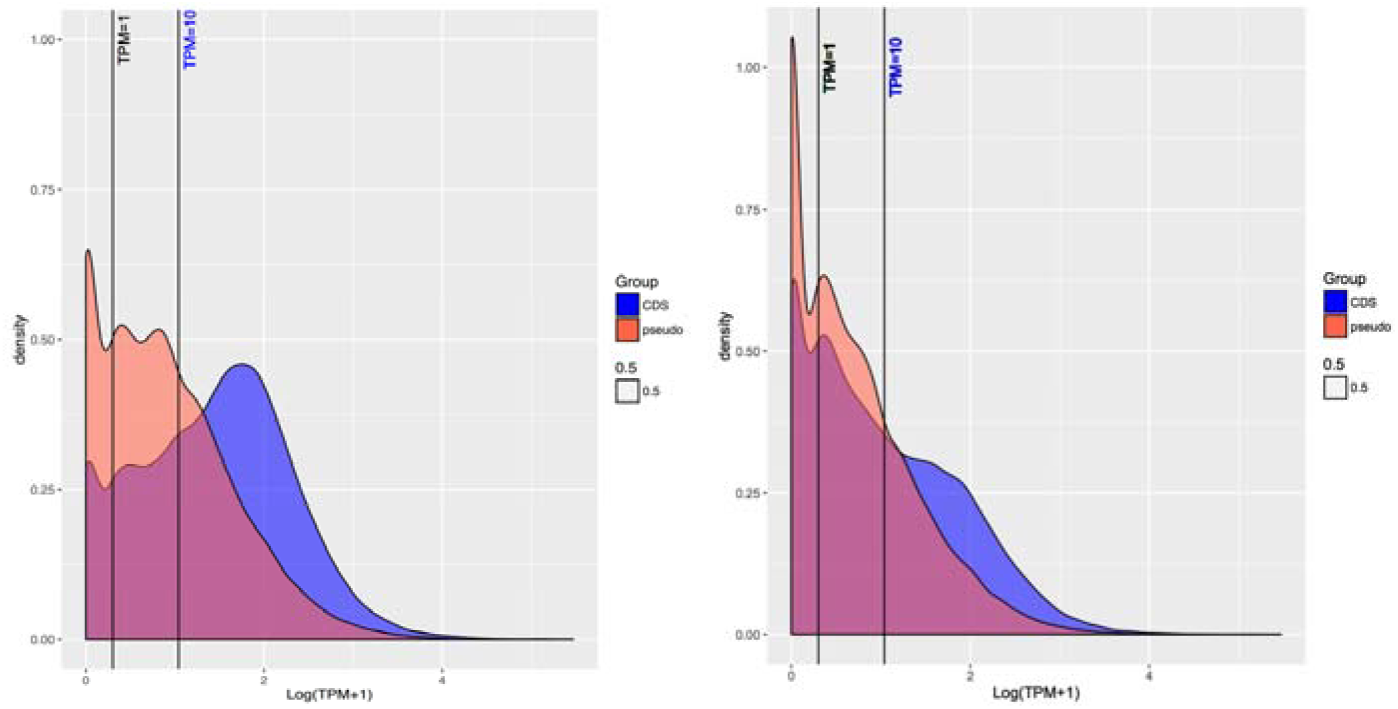
Density plot of expression (Log (Transcripts per Million + 1)) showing transcription for all three cell-free culture conditions, grouped by CDS (blue) or pseudogene (red). Lines represents TPM = 1 and TPM = 10 that represents two different minimum thresholds to be considered as activity. Figure 1B is sense transcription only, and Figure 1C displays all transcription. CDS can be seen to show greater levels of higher expression levels than pseudogenes (red). Overlapping low-expression CDS and pseudogenes highlights the difficulty in identifying pseudogenes using transcription levels.

Across all conditions, mean sense CDS expression (TPM=245.33) is significantly greater than that of pseudogenes (TPM=60.96; Mann-Whitney U Test; W = 46797000, p-value < 2.2e-16). Mean antisense CDS expression (TPM=24.74) is also significantly greater than that for pseudogenes (TPM=13.64; W = 37651000, p-value < 2.2e-16). Actively expressed pseudogenes (TPM≥10) are enriched for organonitrogen compound metabolic processes (GO:1901564; 51 pseudogenes; 31 CDS; p<0.0001). Putatively expressed pseudogenes (TPM≥1 and TPM<10) tend to be involved in transposase activity (GO:0006313; 52 pseudogenes; 14 CDS; p<0.0001).

Differential expression analysis suggests that, of the actively transcribed genes, 938 CDS and 219 pseudogenes are being differentially expressed between either LLP or LSP when compared to ELP growth (FDR ≤ 0.05). In terms of antisense expression: 219 CDS and 106 pseudogenes showed differential antisense expression between timepoints. DE pseudogenes are involved with nucleic acid and organic cyclic compound binding activities (GO:0003677 and GO:0097159).

Two related examples of specific gene degradation putatively associated with positive or negative selection are the type-III secretion and motility systems. *S. glossinidius* has two broad regions that encode for flagella-related proteins; one intact flagellum region (Flagellum Region 1), and one is undergoing significant degradation (Flagellum Region 2, Figure 2), and three T3SS regions (SSR1-3), all largely intact. Coding flagella genes were significantly more likely exhibit sense expression (chi-sq=31.06; p<0.0001) and more likely to be differentially expressed than pseudogenised flagella genes in Flagellum Region 2 (chi-sq=35.27; p<0.0001; Figure 2). The intact flagellum region is up-regulated in late log phase and down regulated in early log phase growth relative to average expression. SSR-1 is up-regulated in early log phase and down-regulated in late stationary phase. SSRs 2 and 3 are down-regulated in early growth and up-regulated in late log and late stationary phase, respectively (Figure 2).

**Figure 2:**
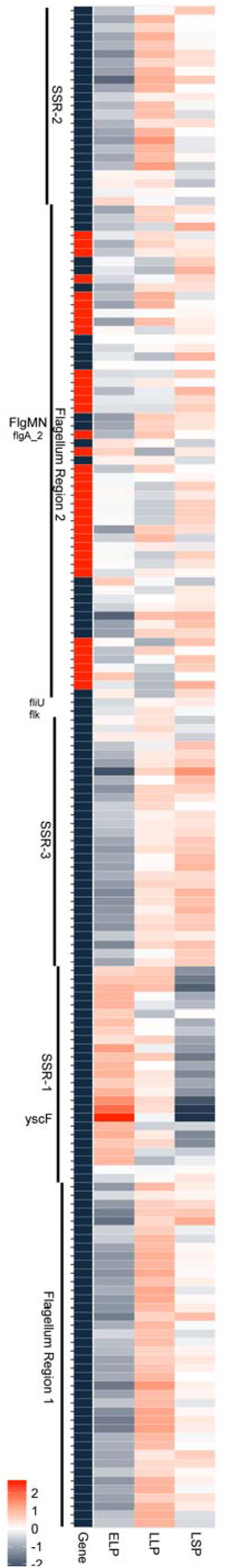
*Sodalis glossinidius* flagellum and *Sodalis* Symbiosis Region expression, summarized by general ‘region’ (left bar). Two genes are not covered by general ‘region’ bars are fliU (top) and flk (bottom). CDS (blue) and pseudogenes (red) are displayed as the first coloured column. Early log phase (ELP), late log phase (LLP) and late stationary phase (LSP) expression is displayed as a heatmap of Log Fold Change relative to average expression, where red signifies up-regulation and blue represents down-regulation.

Figure 3 displays volcano plots of log Fold Change against negative log FDR for Late Log Phase (Figure 3A), and for Late Stationary Phase (Figure 3B), versus Early Log Phase. Each shows that some pseudogenes are highly likely to be differentially expressed between conditions.

**Figure 3:**
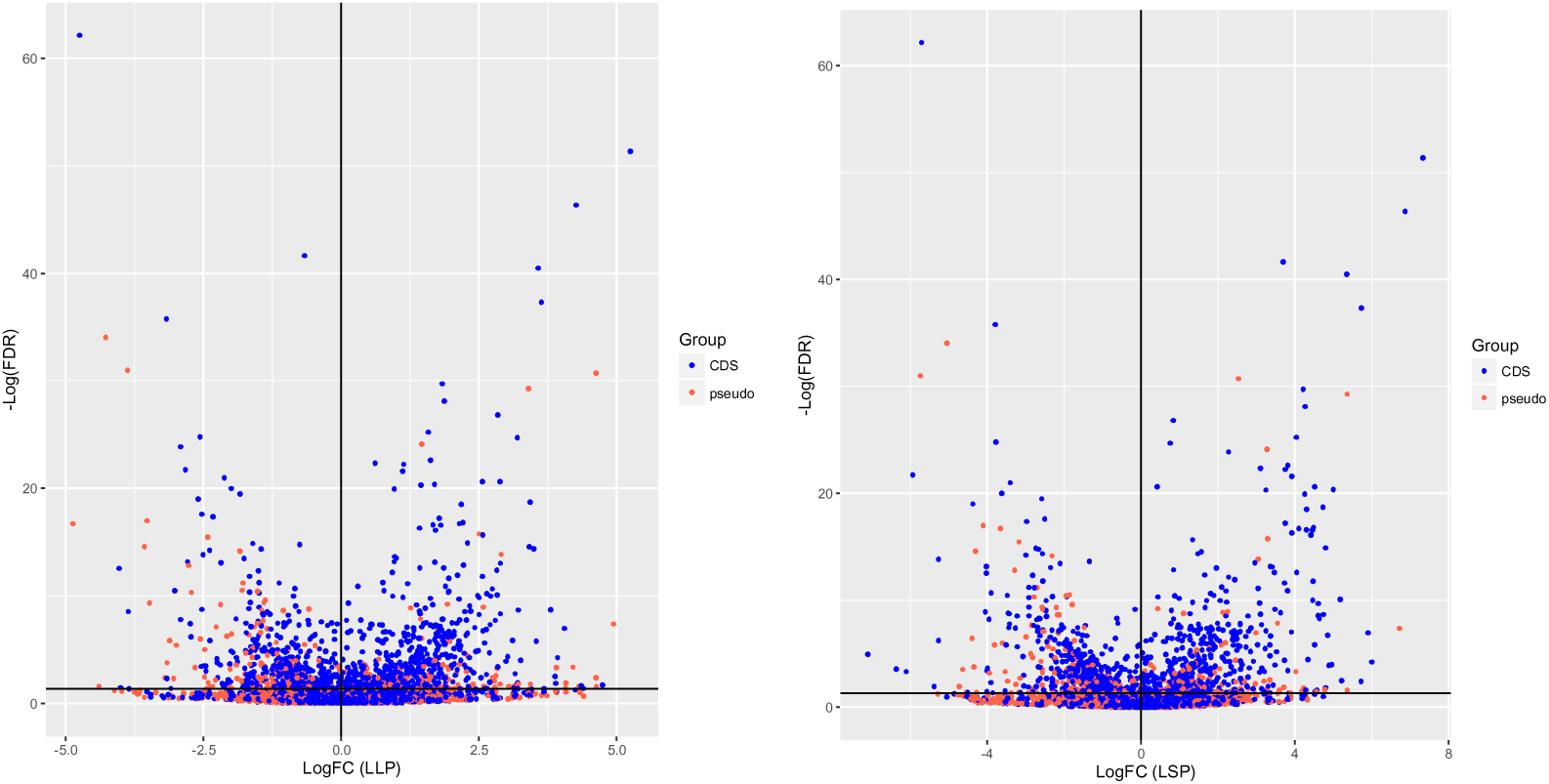
−Log_2_FC differential expression plotted against negative logFDR (False Discovery Rate), determined from EdgeR. Left pane (A) shows Late Log Phase and Right Pane (B) shows Late Stationary Phase transcription. Whilst CDS (blue) represent the majority of differentially expressed genes, some pseudogenes (red) are significantly differentially expressed in either condition.

### SgGMMB4 Proteome

To support pseudogene annotation, and to assess whether transcription may relate to translation we performed proteomic analysis from pooled bacterial cells from all conditions in the cell-free growth experiment. The PROKKA gene models and the 6-frame translations comprised a total of 5,625 and 6,769 proteins respectively. With our MS/MS search on the combined search database, we identified 1,503 peptides attributable to the PROKKA annotation. In two instances, an alternative start codon in the annotation (GTG/Val) was manually changed to ATG to encode a methionine residue, in order to match. We identified 34 pseudogenes from previous annotations that corresponded to the presence of a protein product in our data (Supplementary 2 and 3). Seven pseudogenisation events that split ancestral orthologues into two or more ORFs were verified by observation of peptides from a protein product of one of the resultant ORFs. A further 27 peptides were detected, corresponding to (single-ORF) pseudogene predictions from the original annotation. These were re-classified as CDS and are detailed in Supplementary Data 3. It should be noted that these identified proteins are the *representative proteins* from each protein group as reported by ProteoAnnotator. ProteoAnnotator reports one representative protein from each protein ambiguity group (consisting of one or more proteins) formed due to sharing the same set or subset of peptide identifications. This strategy avoids double counting of proteins with no independent evidence. We further computed the Exponentially Modified Protein Abundance Index (emPAI) value as a semi-quantitative measure of protein abundance for the representative proteins using the mzidLibrary (Supplementary Figure 1) (37).

### SgGMMB4 Methylome and Codon Usage

To assess whether methylation patterns differed between intact CDS and pseudogenes, we used the ability of PacBio SMRT sequencing to detect epigenetic modifications, including (for example), 6mA, 4mC or 5mC, by comparing the sequencing profiles (specifically comparing interpulse duration) between native DNA and PCR-amplified DNA (38). 24,869/29,832 (83.4%) of 5-GATC-3 motifs in the SgGMMB4 chromosome are predicted to be 6-Adenine methylated. No other epigenetic modifications or underlying motifs were detected. In CDS and pseudogenes: CDS display a significantly higher frequency of methylation (4.68 per gene) than pseudogenes (2.54 per pseudogene; Kruskal-Wallis chi-squared = 382.61, df = 2, p-value < 2.2e-16). Mean CDS length is larger than pseudogenes (714 bp vs 415 bp), and although CDS and pseudogenes do not significantly differ in their underlying mean GC-content (CDS=55.48%, pseudogenes=55.71%), there are therefore fewer 5-GATC-3 sites within pseudogenes (mean 3 per pseudogene) than in coding sequences (mean 5.6 per CDS; Kruskal-Wallis chi-squared = 428.82, df=3, p-value < 2.2e-16; Supplementary Data 6). There is an increased frequency of both GAT (increase of 2.7 codons per 1000) and ATC (increase of 2.2 codons per 1000) codons in CDS vs. pseudogenes (Supplementary Figure 2).

### Comparative Genomics to other Sodalis isolates

Comparing SNP rates and pseudogene carriage between multiple genomes of *Sodalis* isolated from different tsetse hosts could reveal whether pseudogenes are being deleted at different rates, or if there is relaxed selective pressure acting on pseudogenes. ROARY pan genome analysis of the six Illumina sequenced isolates compared to our PacBio-sequenced annotation assigned 3,183 CDS and 2,301 pseudogenes to either the core genome (all seven genomes), soft core (2 to 6 genomes) or cloud (one genome). ROARY suggests that there are 2,729 core CDS, 358 soft core CDS, and 184 cloud CDS. 1,796 pseudogenes are assigned to the core genome, which represents ~40% of the overall core genome, despite the phylogenetic distance between hosts. There are an additional 280 soft core and 137 cloud pseudogenes (Figure 4B). Low SNP rates and stable pseudogene carriage, indicated by high numbers of pseudogenes contributing to the *Sodalis* core genome, confirms the suggestion of a recent association of *Sodalis* with the tsetse hosts, and that *Sodalis* developed an association prior to tsetse diversification. This may also imply that either: pseudogenes may not be under relaxed selective pressure, due to the low rate of SNP accumulation; or that there has not been sufficient time since the association for SNP to accumulate. Core single nucleotide polymorphisms (SNP) are not targeted towards pseudogenes: Core single-nucleotide polymorphism using SNIPPY suggests there are 474 core SNP loci in pseudogenes, 814 in intact CDS and 540 intergenic core SNP. Enriched functions of core pseudogenes are transporters (GO:0005215; 144 pseudogenes) and oxidoreductase activity (GO:0016491; 92 pseudogenes). Unique pseudogenes to SgGMMB4 are enriched for transposase (GO:0004803; insertion sequences and phage activity).

**Figure 4:**
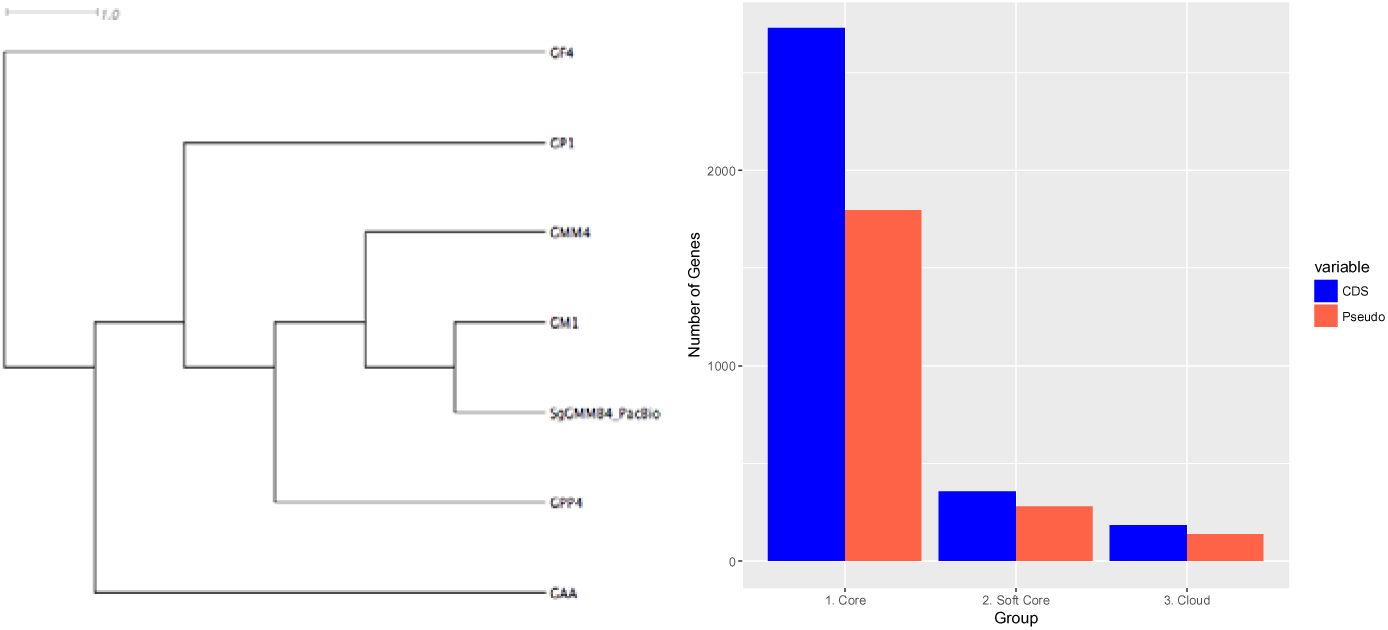
**Pan-genome analysis of** CDS (blue) and pseudogenes (red) from genomes of six *S. glossinidius* isolated derived from four different tsetse species. 2A: GUBBINS-derived whole genome alignment tree showing relatedness between sequenced strains; 2B: Plot representing the number of CDS/Pseudogenes in the ROARY-derived Pan-Genome. Core = 7 genomes; Soft-Core = 2-6 Genomes. Cloud = 1 genome. GAA = *Glossina austeni*; GF = *Glossina fuscipes*; GM(M) = *Glossina morsitans (morsitans)*; GP(P) = *Glossina palpalis (palpalis)*.

Functional corrections of pseudogenes could be important for ecological contribution or evolutionary trajectory of pseudogenes. To test for RNA-based (functional) corrections of genomic SNPs at the transcriptional level, a SNIPPY analysis of the RNA-seq data was performed. A single base insertion in a C(8) homopolymeric tract corrects a frameshift in a putative proline/betaine transporter (Base position 869745, SGGMMB4_01173-01174/SG0498). Neither of the proP2/3 ORFs have been annotated as pseudogenes in this analysis, although no peptide has been detected for either ORF. A second insertion was found to correct for a frameshift in a putative transposase (C(9)>C(10); SGGMMB4_02124/SG0946). Additionally, but not affecting pseudogenes, a single base insertion (G>GG) was found upstream of the annotation for the CDS SlyA_2, which is a transcriptional regulator (SGGMMB4_03264/SG1443). The SNP putatively either alters the region immediately upstream of the 5′-end or extends and alters the first 22 amino acids if re-annotating using an alternative start codon. A BLASTP search of the sequence suggests the current annotation for SlyA_2 is very similar to *SlyA/MarR* orthologues in other endosymbionts (Query coverage ≥98%; e ≤ 4e75). Each of these SNPs was only identified in the RNA-seq data and neither in the Illumina nor PacBio DNA sequencing data.

## Discussion

It seems logical to consider pseudogenes as potentially maintaining a function until their association with transcriptional processes has been silenced. This is particularly pertinent in the case of Secondary (S) symbionts with high proportions of pseudogenes, like *Sodalis*, which are presumed to be evolving towards an obligate association with their host. In secondary symbiosis, the current opinion is that degeneration can largely be attributed to small, vertically transmitted, populations with little diversity reducing the ability of the organism to purge deleterious mutations (i.e. leading to the generation and persistence of pseudogenes) (1, 39, 40).

### Pseudogenes can harbour residual sense and antisense transcription

We have shown that bacterial pseudogenes can be both actively transcribed and dynamically regulated during growth. Expressed genes hypothesized to be under negative selection (i.e. annotations that were significantly more likely to be present in pseudogenes than CDS) were likely to have functions related to transmembrane transport (particularly Major Facilitator Superfamily (MFS) transporters), glycerol metabolism and transposition. MFS pseudogenes permeate genome annotations: A search for “MFS pseudogene” in Ensembl bacteria shows 1,984 genes at the time of writing, across many genera including *Escherichia, Fransicella* and *Pseudomonas* (41). Genes linked to substrate transport (such as MFS transporters) and metabolism are commonly found to be degraded in insect symbiont genomes, linked to their restricted diets (42). Because positive selection is intrinsically linked to substrate bioavailability – as environmental conditions change (such as a lack of nutrients or the provision of a processed metabolite directly to the symbiont) – relaxed selection leads to the accumulation of deleterious mutations and pseudogenisation of redundant genes. Nevertheless, residual expression from these may be indicative of a recent switch in environment, where mutation at the genomic level could outpace overarching transcriptional control, in so much that transcription may be positively selected for due to links to other, maintained, mechanisms.

We have highlighted the related flagella and Type-III secretion / Symbiosis Regions as examples of regions undergoing clear patterns of pseudogenisation (i.e. pseudogenes are clustered with few exceptions, e.g. FlgMN; Figure 2). It is clear that coding genes associated with motility and T3SS apparatus were more likely to be differentially expressed than pseudogenes, and thus the canonical definition of pseudogenes may hold true in these cases. The FlgMN operon remains intact, despite residing within the degrading flagellum region 2, and is differentially expressed in line with Flagellum Region 1 (Supplementary Data 5). Additionally, the only flagellum/T3SS-related pseudogene exhibiting differential expression was flgA_2, which is the only other gene in the FlgMN operon, suggesting that pseudogene differential expression could be a factor of co-regulation of intact genes, or as a result of polycistronic regulation across operons. One of the most differentially expressed pseudogenes, without a hypothetical annotation, is trehalase, an enzyme involved in the breakdown of trehalose – a sugar commonly found in insect haemolymph (43). Amongst a number of hypotheses as to the differential expression of trehelase in cell free culture (which lacks trehalose as a constituent) are implications of residual global control of metabolic processes beyond single sugars. Given the importance of this sugar in insect systems, it would be interesting to further test this pseudogene for residual function.

### Pseudogenes are difficult to define

By using stranded RNA sequencing, and comparing transcription between *Sodalis* CDS and pseudogenes, we have shown that intact CDS show significantly greater mean levels of transcription than pseudogenes, but there remains a proportion of CDS with little or no expression – any of which could be non-functional and mis-annotated. It should be noted that this experiment relied on cell-free culture; CDS may therefore not be expressed due to their functional redundancy in cell-free culture, and follow-up experiments may be required in further media types, or in insect-cell co-culture, to fully ascertain the *Sodalis* transcriptional repertoire. Similarly, however, there remain pseudogenes with residual activity, going against the classical definition of a pseudogene, and it is clear, therefore, that problems remain with the identification and annotation of pseudogenes. We and others have identified novel genes, including genes potentially important in regulating flagellum and/or Type III secretion machinery (*HilA*) or in Quorum Sensing (*SlyA/MarR*) (44). Simply defining pseudogenes using any individual genomic assay is difficult: ORFs may be shortened by frameshift mutations, yet may retain functional domains and appropriate transcriptional architecture. Coding sequences are generally characterized following a set of canonical rules of gene structure: the presence of an open reading frame (ORF), a promoter and ribosomal binding site (RBS); a methionine (or, occasionally, alternative) start-codon and a stop-codon. Similarly, pseudogenes were predicted wherever such rules break down: in the case of *S. glossinidius*, pseudogenes were predicted where <50 % of the functional homologue remained intact. Although studies at the single cell level in *E. coli* (45), and in some conditions at the population level in *Clostridium* (46), suggest that levels of mRNA and protein can remain uncorrelated and be regulated independently of one another, our data suggest that a balance may exist between mRNA transcript and protein abundance, as a semi-quantitative measure of peptide abundance correlates with sense expression. It is likely that each tier of control (i.e. at the DNA, transcription and translation levels), that each may act on another – for instance sRNA may impact mRNA levels, or protein interactions may regulate transcription. There remains a range of bacterial transcriptional processes still to fully characterize, including 5′-UTRs, alternative promoters or alternative transcriptional start and stop sites, and further experiments using techniques such as terminal endonuclease linked RNA-seq, which has been employed in similar experiments in *Salmonella enterica* serovar Typhimurium (47), would shed further light on the transcriptional landscape of this bacterium. It is important to stress that, in this study, we have defined pseudogenes specifically where peptides have *not* been detected; Given the difficulty for proteomics analysis to detect membrane proteins (48), further work may well reveal the presence of hitherto undiscovered peptides and thus that some of these differentially expressed pseudogenes may be mis-annotated and further explain the presence of residual transcription.

### Transcriptional- and post-transcriptional pseudogene control mechanisms remain to be ascertained

Given its dual role in mismatch repair and the regulation of gene expression, Dam-mediated methylation of 5-GATC-3 motifs in bacterial genomes represents a potentially important factor to investigate. While Pacific Biosciences sequencing allowed for the examination of methylation status by comparing modified to unmodified DNA, the potential role methylation might play in pseudogene control remains difficult to ascertain: Pseudogenes displayed a significantly decreased rate of 6mA methylation, when compared to CDS, probably due to the tendency for pseudogenes to have fewer 5-GATC-3 methylation motifs at the genetic level (because pseudogenes are smaller than CDS). Dam-mediated methylation is predicted to post-transcriptionally regulate gene expression by altering the affinity of proteins for DNA, such as at the origin of replication (oriC) (49). In *S. enterica* serovar Typhimurium, Adenine methylation has been implicated in regulating quorum sensing derived virulence factors and as such Dam inhibitors or Dam-silenced pathogens have been studied for their antimicrobial or vaccine potential, respectively (50). Adenine methylation has also been implicated in protecting symbionts from heat-stress (51).

### Pseudogene abundance is stable between Sodalis genomes

Given that we expect *Sodalis* to be routinely undergoing population bottlenecks through vertical transmission of their host, we could expect genetic drift to be acting on genes under little selective pressure, increasing SNP and/or pseudogenisation rates, or even driving their deletion. As accessory genomes diverge prior to SNP arising in the core genome (52), examining the *Sodalis* pan-genome derived from *S. glossinidius* species from multiple tsetse hosts, enabled us to examine pseudogene stability. The high number of pseudogenes in the *Sodalis* core genome suggests pseudogenes are stable across *Sodalis* strains infecting different tsetse species – in line with the suggestion that *Sodalis* shares an evolutionarily recent association with its tsetse host due to a lack of concordant phylogeny, in contrast to that of *Wigglesworthia*, its primary symbiont (53). Lawrence *et al* (2001) suggested that intracellular lifestyle promotes protection from bacteriophages and insertion sequences, reducing recombination and promoting pseudogene persistence (54). *S. glossinidius* GMBB4 has 44 chromosomal prophage elements, and an active circular phage (pSG3). Furthermore, sexual transmission of *Sodalis* from father to mother prior to vertical transmission has been reported, effectively increasing diversity and rates of recombination (55). Assessing *Sodalis* genetic variability across wild populations of tsetse, rather than lab-reared tsetse will be essential in understanding pseudogene persistence. Similarly, further research into expression levels of both coding and pseudogene orthologues may reveal further insight into transcriptional co-regulation – and gene redundancy linked to reductive evolution (56). Kuo and Ochman have previously suggested that *Salmonella* pseudogenes may lack sufficient negative pressure for deletion (11). In models of cyanobacterial genomes, increased resource levels and decreased mortality have been suggested to select for slower reproduction and streamlined genomes (57). Experimental evolution experiments in *Methylobacterium* have shown that accessory gene deletion confers a direct fitness benefit under selective environments, rather than the associated benefit of the reduced fitness costs of maintaining a shorter genome in its own right (58), and similar experiments would be interesting to perform in this system.

Defining which associations constitute significant function for which positive pressure ensures their persistence is difficult: associations with promoters, transcription factors or *cis/trans*-acting transcriptional regulators could all select for pseudogene retention in the genome, and reduce the likelihood of full deletion. An increasing number of bacterial small RNAs have been identified through transcriptomic analyses, including in *Streptococcus* (59) and *Borrelia* (60). Pseudogene-derived antisense RNA may be involved in the complex interactions between genome, transcriptome and proteome: sRNAs have been implicated in gene regulation of multiple target genes through processes such as translational inhibition and activation, or transcript stability (61). *Hfq*, a crucial chaperone involved in bacterial sRNA processing, is maintained in *Sodalis* (SGGMMB4_00878). Therefore, further studies into the roles of *Sodalis* sRNAs – including pseudogene-derived sRNAs and the role of *Hfq* or other chaperones – will be critical for full understanding of the complexity of gene regulation in this degrading bacterial genome. This is in line with previous work using tiling arrays in *Mycoplasma pneumonia*, wherein frequent antisense and non-coding transcripts were identified in a degrading bacterial genome (62). Pseudogene-derived transcripts (such as antisense small RNAs derived from pseudogenes) could act as regulators for orthologues elsewhere in the genome (63) and such an association may reduce the selective pressure towards their deletion. As this study relied on RNA-shearing based library preparation, it would be interesting to follow up this study with full-length third-generation RNA sequencing (i.e. using Pacific Biosciences cDNA (Isoseq) or direct RNA sequencing using Oxford Nanopore technologies) to elucidate the sequences of full-length mRNA, including primary, polycistronic transcripts, which would further enhance our knowledge as to how pseudogenes continue to contribute to overall transcription and its control despite ongoing genomic degradation.

## Conclusions

The primary goal of this study was to establish whether bacterial pseudogenes remain active despite genomic degradation, using *Sodalis* as a model given the number of putatively inactivated, functionless genes persisting in its genome. We sought to combine DNA sequencing, stranded RNA-sequencing and proteomic analysis to fully describe the Sodalis transcriptional and translational landscape with a view to better understand the evolution and functional control of bacterial pseudogenes and the process of endosymbiont genome degradation.

We have revealed that whilst transitioning from a free-living to symbiotic status, *Sodalis* pseudogenes are often transcribed, but at a significantly lower level than intact CDS. Some pseudogenes even remain under active transcriptional control, exhibiting differential expression throughout growth, however proteomic analysis suggests they ultimately do not contribute to the protein content of the cell. The lack of some expression from intact CDS and pseudogenes underpins the difficulty in pseudogene identification – especially in cell-free culture where the correct conditions for their expression may be lacking. That a combination of sense and antisense transcription of pseudogenes persists implies a role of pseudogene transcription in control mechanisms: e.g. *cis/trans* small RNA transcriptional control, and could even be playing a role in wide-reaching mechanisms such as host-symbiont interaction, or symbiont-symbiont interaction. The persistence of pseudogenes in the *Sodalis* pangenome implies that the maintenance of function of degraded genes may outweigh any deleterious effects, or that there exists a mechanism by which such deleterious effects are mitigated. Given the proximity of *Sodalis* to medically important parasites and other bacteria within the tsetse host, further study on these mechanisms is of interest for identifying novel therapeutic interventions.

## Materials and Methods

### DNA Sequencing

For PacBio sequencing: *Sodalis glossinidius* strain GMMB4 (SgGMMB4) was isolated from *Glossina morsitans morsitans* (Westwood) from the Langford derived long-term colony maintained at the University of Edinburgh in 2005. Six further *S. glossinidius* isolates were cultured from lab-based tsetse for Illumina sequencing: GP1 and GPP4 were isolated from *Glossina palpalis palpalis*; GAA from *G. austeni*; GF4 from *G.fuscipes*; GM1 and GMM4 from *G. morsitans* as previously described (64).

Bacteria were recovered from -80 °C storage by incubation at 25 °C on columbia agar plates supplemented with 10 *%* defibrinated horse blood (TCS Biosciences) in microaerophilic conditions (~5-12% CO_2_ CampyGen, Oxoid, UK). An individual colony was picked and grown to late stationary phase in cell-free culture medium at 25 °C in Schneiders Insect Medium (Sigma, UK) supplemented with 10 % Fetal Calf Serum (Life Technologies, UK). High molecular weight whole genomic DNA (gDNA) was extracted from the subsequent bacterial pellet using the Zymo Research Universal gDNA extraction kit (SgGMMB4) or the Qiagen DNeasy kit (other isolates) according to the manufacturers instructions.

SgGMMB4 gDNA was sequenced on the Pacific Biosciences RS-II instrument (PacBio) at the Centre for Genomic Research at the University of Liverpool on a single SMRTcell using P6-C4 chemistry with no prior size selection. Reads were assembled and contigs polished using HGAP.3 resulting in a polished assembly consisting of a single chromosomal contig and nine further contigs. Comparison of the sequence to the available reference by MUMMER (65) and ACT (66) suggested that two contigs were a result of chimeras derived from pSG2. A further five contigs were repetitive phage-derived sequences. The chromosome was subsequently manually edited to begin at the start of the dnaA gene. The putative protein-coding, ncRNA, and tRNA gene sequences were annotated using PROKKA (v. 1.10) (67). Pseudogenes in this study were initially conservatively annotated where the PROKKA-defined ORFs overlapped with the Belda-annotated pseudogenes or predicted to be pseudogenes by PROKKA (67). In the latter case, PROKKA predicted pseudogenes based on identical annotations in sequential open reading frames (except for hypothetical protein annotations). Sequences matching possible riboswitch domains were predicted using the Denison Riboswitch Detector online webserver (Supplementary Data 4) (68). Additionally, the two available annotations for SgGMM4 (14, 23) were transferred to the PacBio SgGMMB4 scaffold using the RATT software package (32), for comparison and pseudogene prediction. An additional *S. glossinidius* sample (isolated from *Glossina palpalis)* was sequenced and assembled in the same manner using two SMRTcells using P6-C4 chemistry.

Sequencing libraries for the six further isolates of *Sodalis glossinidius* from multiple tsetse species (GAA; GF4; GM1; GMM4; GP1; GPP4) were prepared using a TruSeq library preparation kit (according to the manufacturer’s instructions) and sequenced on a single lane of an Ilumina HiSeq (High-output run; 2x100bp paired end reads) at the Centre for Genomic Research. The Illumina HiSeq data from the six further *S. glossinidius* isolates were initially processed using CASAVA 1.8 to produce FASTQ files. FASTQ data files were trimmed for the presence of Illumina adapter sequences using Cutadapt (v1.2.1), using the –O 3 option (69). The reads were further trimmed using Sickle (v1.200) (https://github.com/najoshi/sickle) with a minimum window quality score of 20. Data were assembled *de novo* with SPADES using default parameters and annotated using PROKKA as previously described. PROKKA-derived GFF annotations were processed through the ROARY pan-genome package to ascertain core and accessory genome coverage (70). Reads were mapped to the SgGMMB4 PacBio reference, and core SNP phylogenies derived using the SNIPPY package (https://github.com/tseemann/snippy).

### Methylome sequencing

In addition to gDNA sequencing and assembly, the PacBio RSII instrument can detect epigenetic modification either in silico or by comparing native DNA to a PCR control. To that end, a Whole Genome Amplified (WGA) control was generated from SgGMMB4 as follows: 1μg gDNA was split into three equal reactions and Whole Genome Amplified using the Qiagen Repli-g Turbo kit according to the manufacturers instructions. These were then pooled and cleaned using a 2:1 ratio of homemade SPRI bead cleanup system analogous to Ampure XP beads (71). The WGA control was sequenced using one SMRTcell in the same way as described previously. Comparison to the native DNA was performed using the Motifs and Modifications module within the SMRT^®^ Analysis Server with a mapping quality cutoff set at QV70 and the subsequent modifications and motifs file filtered for those with a quality ≥ Q50 (P<0.0001).

### RNA sequencing

Individual 10 mL cell-free liquid cultures, as described above, were set up in quintuplets for seventeen time-points at six-hourly intervals. At each time point, the contents of the culture flasks were transferred to 15 mL Falcon tubes, an Optical Density (600nm) measurement taken, and then centrifuged for 10 min at 10 °C. The bacterial pellet was immediately resuspended in 1 mL Trizol reagent (Life Technologies) and total RNA was extracted using Zymo Research DirectZol columns. RNA cleanups were performed using a 2:1 ratio of SPRI beads as described previously. Three timepoints, representing: Early Log (12 hours); Late Log (72 hours) and Late Stationary phase (108 hours) according to the OD measurements were selected and DNase I treated using a Life Technologies DNAfree Turbo kit (according to the manufacturers instructions; (data not shown)).

Ribosomal RNA (rRNA) was depleted using a Ribo-Zero™ bacterial (low-input) rRNA Removal Kit (Epicentre), and individually barcoded, strand-specific Illumina cDNA libraries were prepared using a NEBNext^®^ Ultra™ RNA Library Prep Kit for Illumina. Sequence data was generated using one Illumina MiSeq run with v2 chemistry generating 250 bp paired-end reads. All RNA and cDNA cleanups were performed using SPRI beads as described previously. All raw Illumina sequence Fastq files were trimmed for the presence of adapter sequences using Cutadapt version 1.2 using option –O 3 (69) and quality-trimmed using Sickle version 1.200 (72) with a minimum window quality score of 20. Any reads shorter than 10 bp after trimming were removed. Quality scores for all sequences were assessed using FASTQC v0.9.2 (http://www.bioinformatics.babraham.ac.uk/projects/fastqc). RNA-seq reads were aligned to the PacBio RSII-generated SgGMMB4 scaffold using Bowtie2 v. 2.1.0 (73). The resulting SAM files were converted to BAM and sorted using samtools v.0.1.18-r580 (74)s. For transcript-based annotation, reads were counted against the PROKKA/PacBio annotation using HTSeq version 0.5.3p9 using the –stranded option, to count in both the sense and antisense directions, and using the intersection-nonempty mode (75). EdgeR analysis was implemented using the DEGUST web package (http://degust.erc.monash.edu), which outputs counts per million, logFC and differential expression statistics (Supplementary Data 1). EdgeR counts per million were transformed to TPM according to Wagner *et al* (2012) (76). Further statistical analysis and figure plotting were implemented using R version 3.32. Operon structure was predicted from the RNA-seq data using Rockhopper with the default settings (77). The combined RNAseq sequencing reads were fed through the SNIPPY pipeline (as previously described) to identify potentially correcting SNPs. Gene enrichment analyses were performed in BLAST2GO 5, using Interpro GO IDs (Supplementary Data 1).

### Proteomics

Liquid cultures were grown as previously described for early- mid-log and late-stationary growth phase *S. glossinidius GMMB4*. PBS-washed pellets were suspended in 250 μl of 25 mM ammonium bicarbonate and sonicated, using a Sonics Vibra Cell (Sonics and Materials Inc., Newton, U.S.A.) and 630-0422 probe (250 μl to 10 mL) for a total of 120 joules. The sample was then analysed for protein content, 50 μg was added to 0.05% RapiGest™ (Waters, Manchester) in 25 mM ammonium bicarbonate and shaken at 550 rpm for 10 min at 80°C. The sample was then reduced (addition of 10 μl of 60 mM DTT and incubation at 60 °C for 10 minutes) and alkylated (addition of 10 μl of 180 mM iodoacetamide and incubation at room temperature for 30 minutes in the dark). Trypsin (Promega U.K. Ltd., Southampton, proteomics grade) was reconstituted in 50 mM acetic acid to a concentration of 0.2 μg/μl and 10 μL added to the sample followed by overnight incubation at 37 °C. The digestion was terminated and RapiGest™ removed by acidification (1 μl of TFA and incubation at 37 °C for 45 min) and centrifugation (15,000 x g for 15 min). To check for complete digestion each sample was analysed pre- and post-acidification by SDS-PAGE.

For LC-MS/MS analysis, a 2 μl (1 μg) injection was analysed using an Ultimate 3000 RSLC™ nano system (Thermo Scientific, Hemel Hempstead) coupled to a QExactiveHF™ mass spectrometer (Thermo Scientific). The sample was loaded onto the trapping column (Thermo Scientific, PepMap100, C18, 300 μm X 5 mm), using partial loop injection, for seven minutes at a flow rate of 4 μl/min with 0.1 % (v/v) FA. The sample was resolved on the analytical column (Easy-Spray C18 75 μm x 500 mm 2 μm column) using a gradient of 97% A (0.1% formic acid) 3% B (99.9% ACN 0.1 % formic acid) to 70% A 30 %B over 120 minutes at a flow rate of 300 nl min^−1^. The data-dependent program used for data acquisition consisted of a 60,000 resolution full-scan MS scan (AGC set to 3e6 ions with a maximum fill time of 100 ms) the 18 most abundant peaks were selected for MS/MS using a 30,000 resolution scan (AGC set to 1e5 ions with a maximum fill time of 45 ms) with an ion selection window of 1.2 m/z and a normalised collision energy of 28. To avoid repeated selection of peptides for MSMS the program used a 30 second dynamic exclusion window.

The protein identification for the MS/MS dataset was performed using an open-source software tool – ProteoAnnotator (78). ProteoAnnotator provides an automated pipeline for various interconnected computational steps required for inferring statistically robust identifications. The tool produces a variety of output files compliant with the data standards developed by Proteomics Standard Initiative (79). Mass spectra in form of a MGF (Mascot Generic format) file were provided as input to the tool, along with the search criteria and protein database as described in the sections below. The search parameters for the MS/MS dataset were fixed modification of carbamidomethylation of cysteine and variable modification of oxidation of methionine. A single missed trypsin cleavage was allowed. The product tolerance was set as ±0.5 Da and the precursor tolerance was set as 10 ppm. The protein search database comprised of the gene model predicted by PROKKA, as previously described, plus a six-frame translation of the SgGMMB4 genome. Six-frame translated sequences with length less than eight were excluded from the search database. Decoy sequences were added to the database with a true:decoy ratio of 1:1 to create a final protein database for performing the MS/MS search. For the post-processing of results, we applied a threshold of 5% for both peptide level and protein group level FDRs as described in Ghali *et al* (2004).

## Nucleotide sequence database identifiers

The Pacific Biosciences assembly and annotation have been submitted to the European Nucleotide Archive under accessions LN854557-LN854560. Illumina sequence reads for the six additional *Sodalis* isolates are available under Project accession PRJEB9474 (accessions ERR2036891-ERR2036896). RNAseq data are available under project PRJEB20150.

The mass spectrometry proteomics data have been deposited to the ProteomeXchange Consortium via the PRIDE (80) partner repository with the dataset identifier PXD007068.

## Acknowledgements

This research was funded by a BBSRC project grant BBJ017698/1 to ACD. The Pacific Biosciences RS-II instrument purchase was funded by the BBSRC under grant BB/L014777/1. IG is supported by a Wellcome Trust Seed Award in Science (Ref: 200690/Z/16/Z). Pisut Pongchaikul was supported by a Maihidol studentship. All sequencing was carried out at the Centre for Genomic Research at the University of Liverpool, UK. Thanks also to Dr. Chandan Pal, and the Hinton and Kröger research groups for their support and useful discussions, and to anonymous reviewers whose comments helped us to significantly improve the manuscript.

## Supplementary Data

### Supplementary Data 1

**Differential Expression analysis for Sodalis glossinidius SgGMMB4 (SuppData1.xlsx)**

**Supplementary Figure 1:**
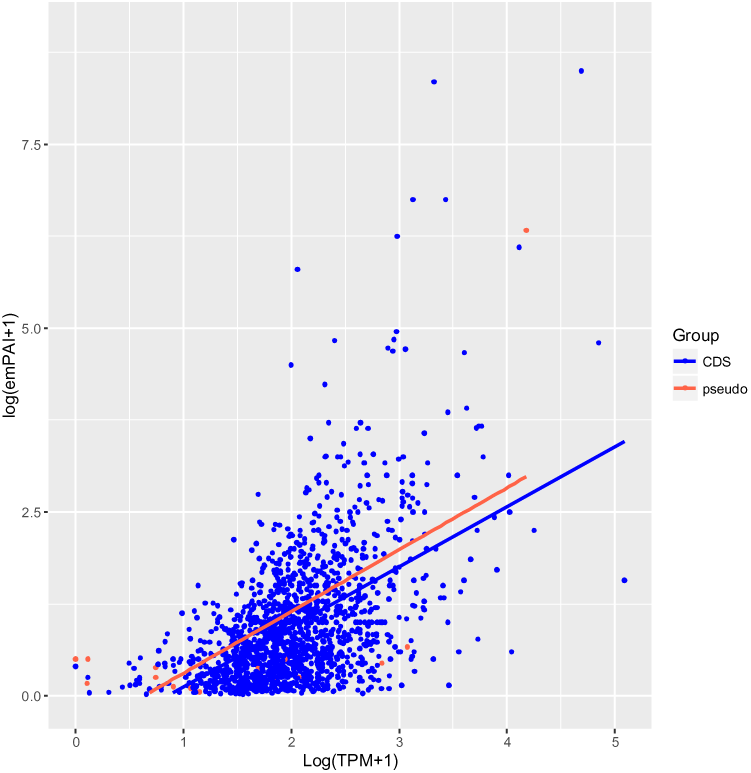
Transcription vs. semi-quantitative protein abundance (for all detected peptides). Line represents linear regression (implemented using ggplot2 in R).

### Supplementary Data 2-4 (SuppData2_3_4.docx)

Supplementary Data 2: Pseudogenised genes with residual active translation

Supplementary Data 3: Table of Single ORF Pseudogenes re-annotated due to detected peptide products

Supplementary Data 4: Riboswitch predictions

### Supplementary Data 5

**Flagellum/Type III Secretion Data (SuppData5.xlsx)**

### Supplementary Data 6

**Summary of Methylation and Codon Usage analysis (SuppData6.xlsx)**

